# Collinearity Analysis of Allotetraploid *Gossypium Tomentosum* and *Gossypium Darwinii*

**DOI:** 10.1101/031104

**Authors:** Fang Liu, Zhong L. Zhou, Chun Y. Wang, Yu H. Wang, Xiao Y. Cai, Xing X. Wang, Kun B. Wang, Zheng S. Zhang

## Abstract

*G. tomentosum* and *G. darwinii* are wild allotetraploid cotton species, characterized with many excellent traits including finer fiber fineness, drought tolerance, Fusarium wilt and Verticillium wilt resistance. Based on construction of F_2_ linkage groups of *G. hirsutum* × *G. tomentosum* and *G. hirsutum* × *G. darwinii,* two genetic linkage maps were compared. As a result we found a total of 7 inversion fragments on chr02, chr05, chr08, chr12, chr14, chr16 and chr25, 3 translocation fragments on chr05, chr14 and chr26. Further comparing the inversion and translocation fragments, we noticed four of seven markers orientation of *G. tomentosum* consistent with *G. hirsutum or G. raimondii,* one of seven inversion markers orientation of *G. darwinii* consistent with *G. hirsutum*; meanwhile one of three translocation marker orientation of *G. tomentosum* consistent with *G. raimondii.* The result indicate, compare *G. darwinii, G. tomentosum* has closer genetic relationship to *G. hirsutum.* This study will play an important role in understanding the genome structure of *G. tomentosum* and *G. darwinii,* and open the doors for further in-depth genome research such as fine mapping, tagging genes of interest from wild relatives and evolutionary study.

## Introduction

Cotton is a natural vegetable white fibrous agricultural product of great economic importance as raw material not only for textile industry but also for a wide variety of uses such as paper industry, home fixtures, medical supplies, chemicals and oil (Ensminger et al. 1990). Cotton (*Gossypium* spp) is one of the most expansively grown species around the globe in tropical and subtropical regions between 36° South latitude and 46° North latitude (Reller et al. 1997) of more than 80 countries(Fryxell 1979; Smith 1995). *Gossypium* genus consists of 50 species (Fryxell 1992; Stewart 1995; Ma et al. 2008), which includes five allotetraploid species (2n=4x=52; (AD)_1_ to (AD)_5_) and 45 diploid species (2n=2x=26, A through G and K) (Fryxell 1979; Stewart 1995; Brubaker et al. 1999; Zhang et al. 2005). The evolution of domesticated cotton involves various steps that can help scientists to have deep insight understanding of polyploidy role in diversification process and better investigation of unique traits, and it can further help geneticists to utilize germplasm from wild relatives. Allopolyploids provide a great tool to understand the evolutionary process of two diploid genomes which evolve simultaneously but different in genome size. Evolution and diversity studies of *Gossypium* genus provide the basic knowledge of morphological diversity of the genus and plant biology which can help in the better utilization of genetic resources (Wendel et al. 2009).

More than 30 genetic maps have already been published in cotton, and most of them are based on interspecific crosses of domesticated tetraploid species namely *G. hirsutum* and *G. barbadense* (Jiang et al. 1998; Zhang et al. 2002; Lacape et al. 2003; Nguyen et al. 2004; Rong et al. 2004; Guo et al. 2007; He et al. 2007; Lacape et al. 2009). The interspecific tetraploid genetic maps are useful in understanding genome structure and exploring the genetics basis of important agronomic characters and also provide the basis for finding new DNA markers for further high density maps (Guo et al. 2007; Zhang et al. 2008b; Yu et al. 2011).

Wild cotton has long been used as a genetic resource to introduce new traits to increase the potential of cotton cultivated species (Stewart 1995). *G. darwinii,* a wild allotetraploid species with (AD)_5_ genome, is closely related to *G. barbadense* but quite different from the cultivated *G. hirsutum,* and it has many excellent traits including finer fiber fineness, drought tolerance, Fusarium wilt and Verticillium wilt resistance. *G. tomentosum* is endemic to Hawaiian Islands, and it has many unique agronomic traits such as insect-pest resistance, drought tolerance, salt tolerance, heat tolerance, nectarilessness and lint color. Many genetic studies like genomic and genetic structure/organization have been conducted on the two cultivated tetraploids, *G. hirsutum* and *G. barbadense*, however very little is known about genomic architecture, gene transfer or introgression of unique traits from other three tetraploids.

In the present study, the SSR genetic maps are developed from two F_2_ populations, (*G*. *hirsutum* × *G. tomentosum*) (Chen et al. 2015) and (*G. hirsutum × G. darwinii*) (Kashif et al. 2015). The linearity relationship is compared between the two genetic maps, which will serve as an indispensable genomic resource for genome structure study, comparative genomic analysis, and fine mapping and map-based gene cloning of important traits.

## Materials and methods

Linkage map were reconstructed reference Chen et, al (2015). and Kashif et, al (2015). SSR markers uniform distribution on different chromosomes were selected and checked after downsizing. JoinMap 4.0 software was used for linkage analysis and map construction. Kosambi mapping function was used to convert recombination frequencies into map distances (centimorgan, cM). Mapchart 2.2 software was used to draw the genetic map.

Linkage map assignment was established by the common markers that were already anchored according to the previous publications (Wang et al. 2012; Li et al. 2014; Li et al. 2015; Zhang et al. 2015). Chromosomal nomenclature was used as mentioned by Guo et el. (2008), i.e. SSR loci anchored on chromosome (Chr.) 1 to 13 were designated to the A sub-genome (At), whereas loci confined to Chr. 14 to 26 were designated to the D sub-genome (Dt).

## Results

Comparing the linear relationship of two genetic linkage maps, we found that most SSR markers present collinearity between the two genetic linkage maps. Meanwhile, part of the non linear relationship appeared on the individual chromosomes between the genetic linkage maps of *G. hirsutum* × *G. tomentosum* and *G. hirsutum* × *G. darwinii,* which include 7 inversion fragments and 3 translocation fragments (Figure 1-13).

**Figure 1.**
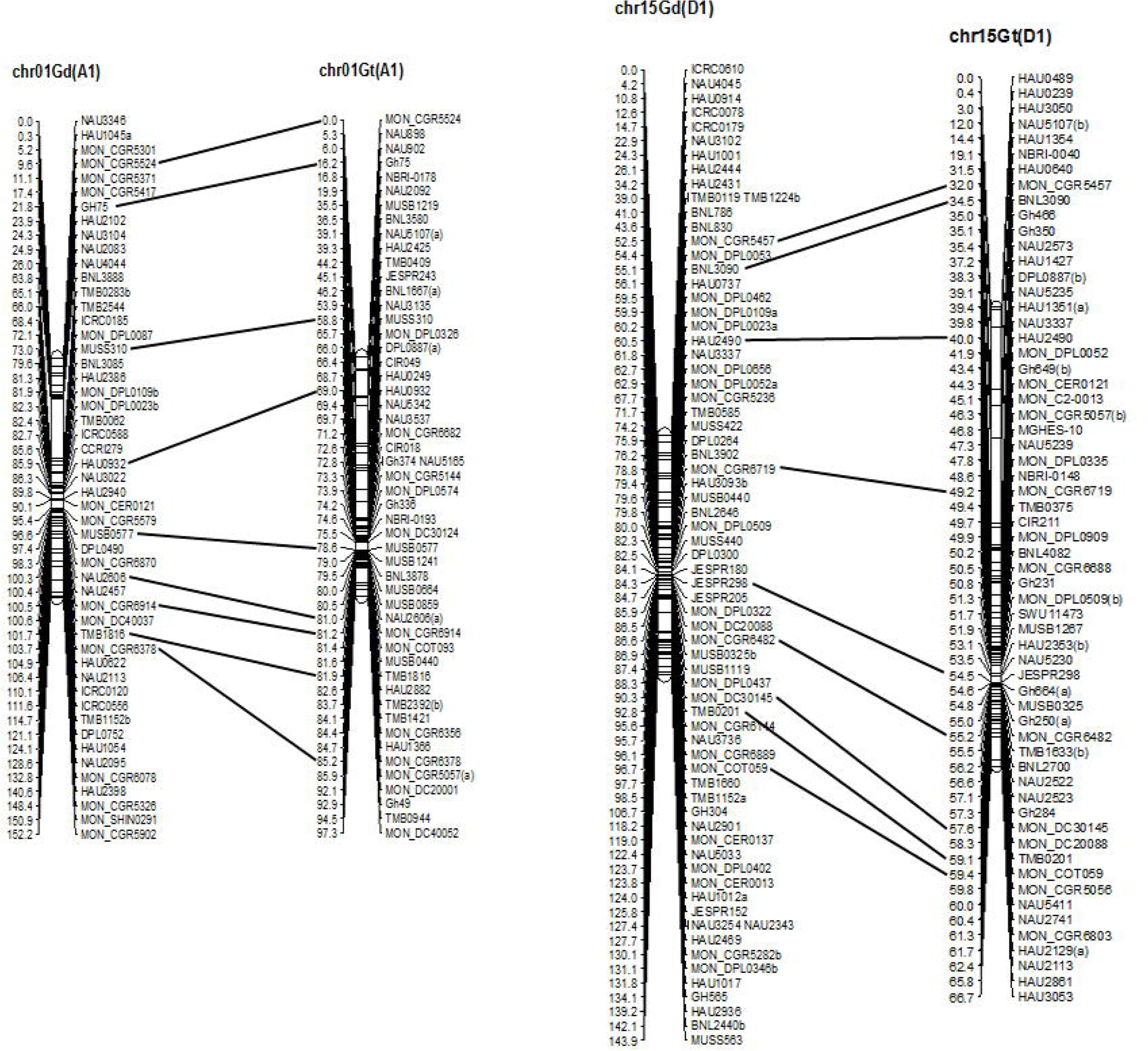

**Figure 2.**
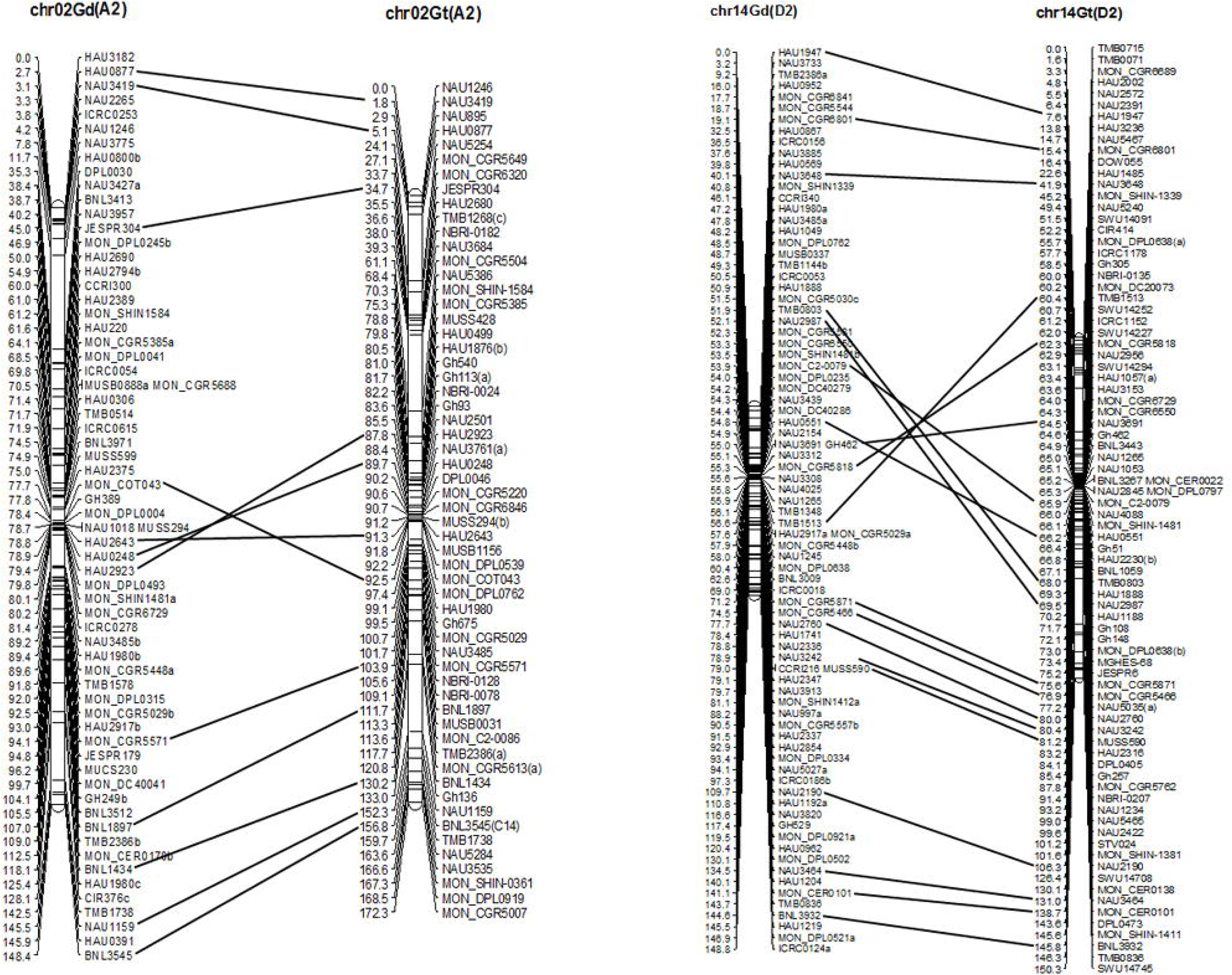

**Figure 3.**
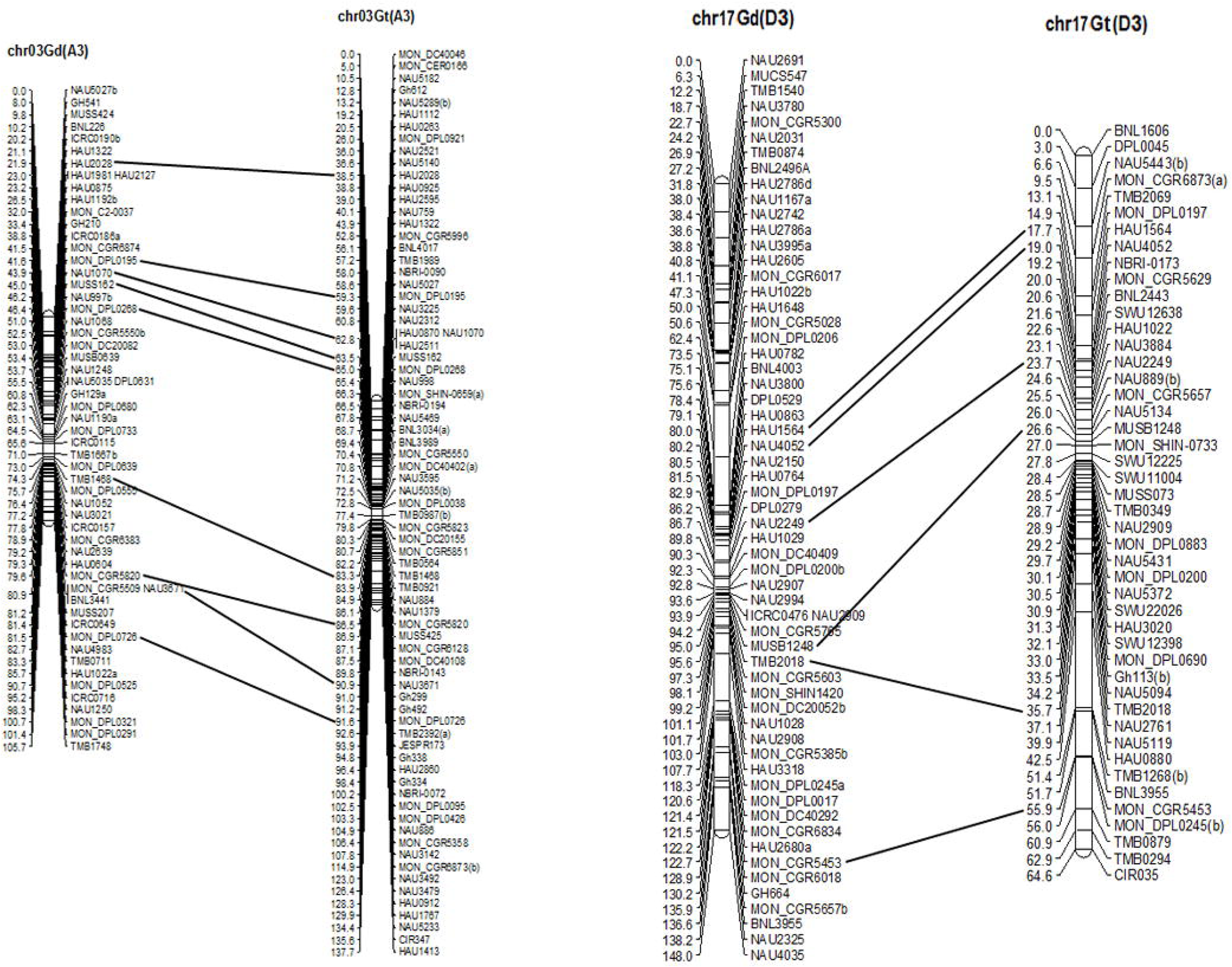

**Figure 4.**
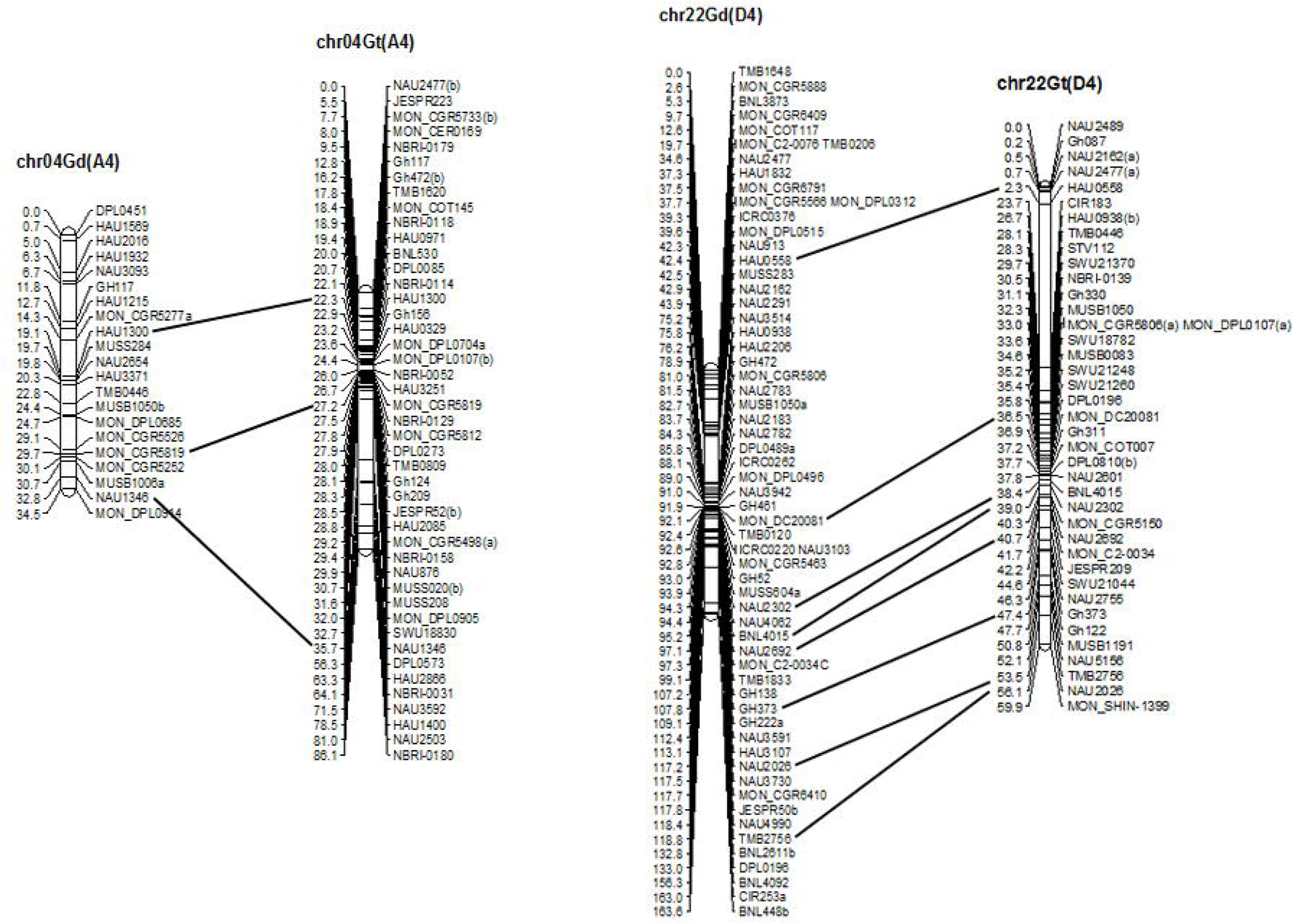

**Figure 5.**
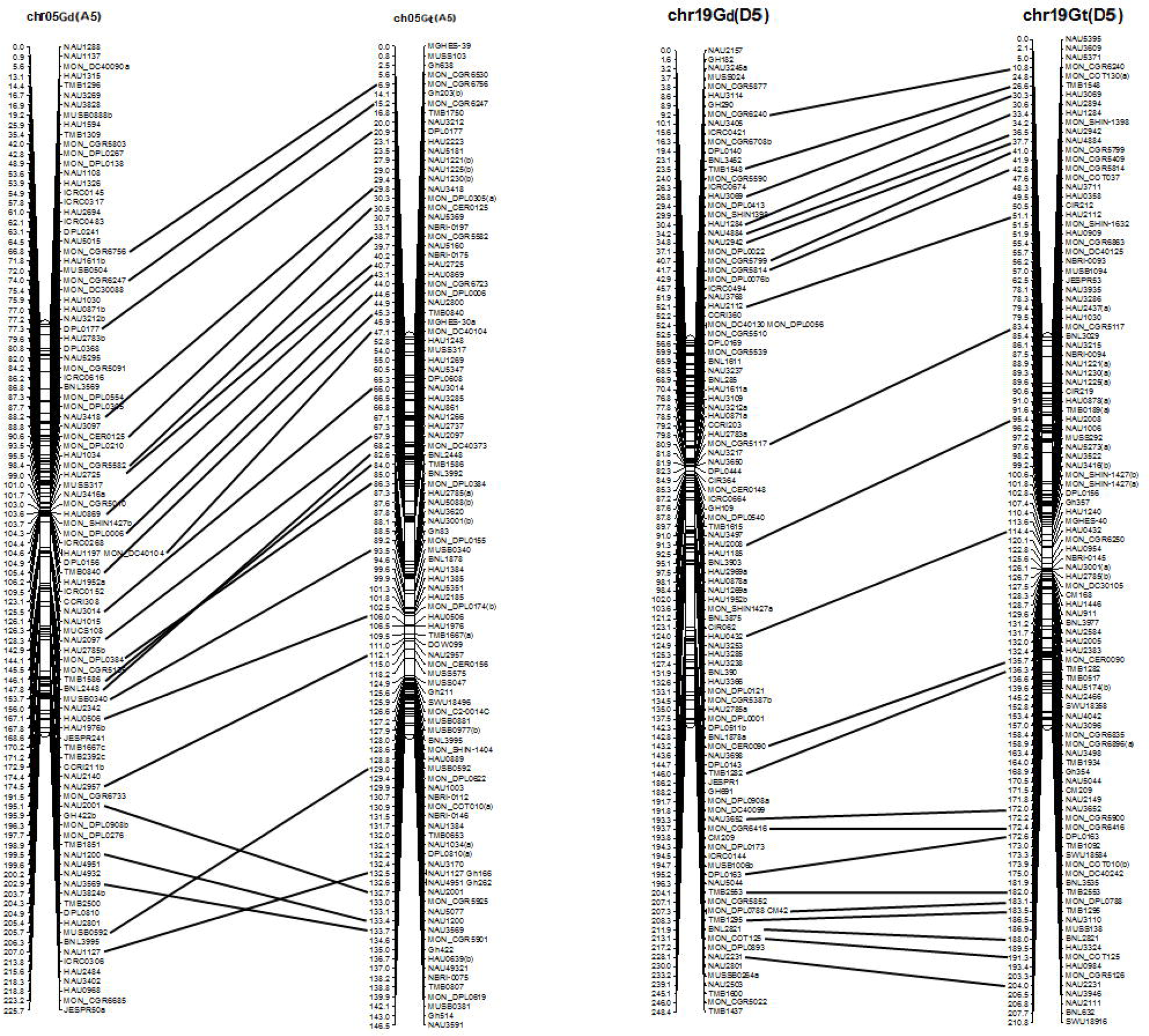

**Figure 6.**
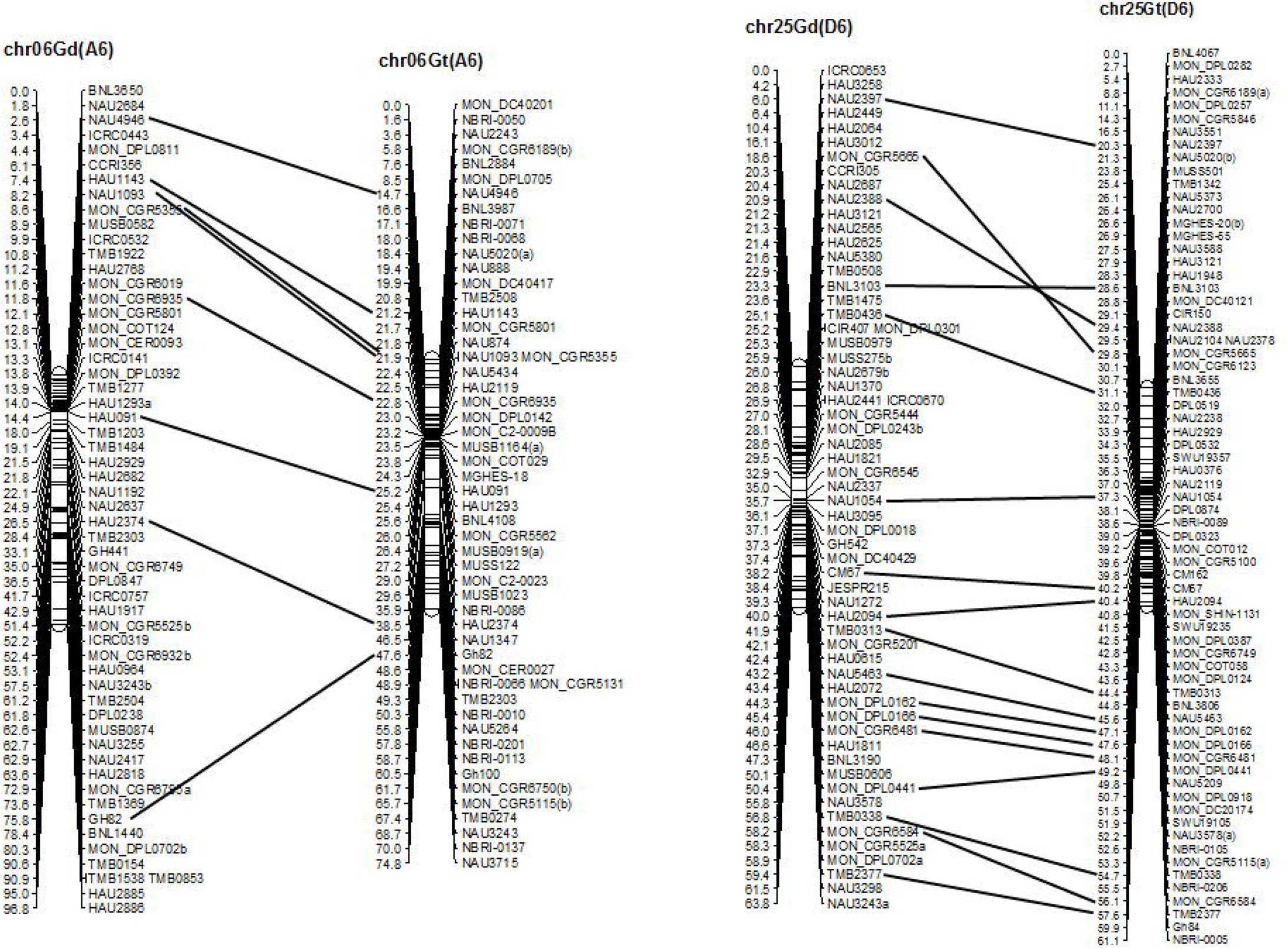

**Figure 7.**
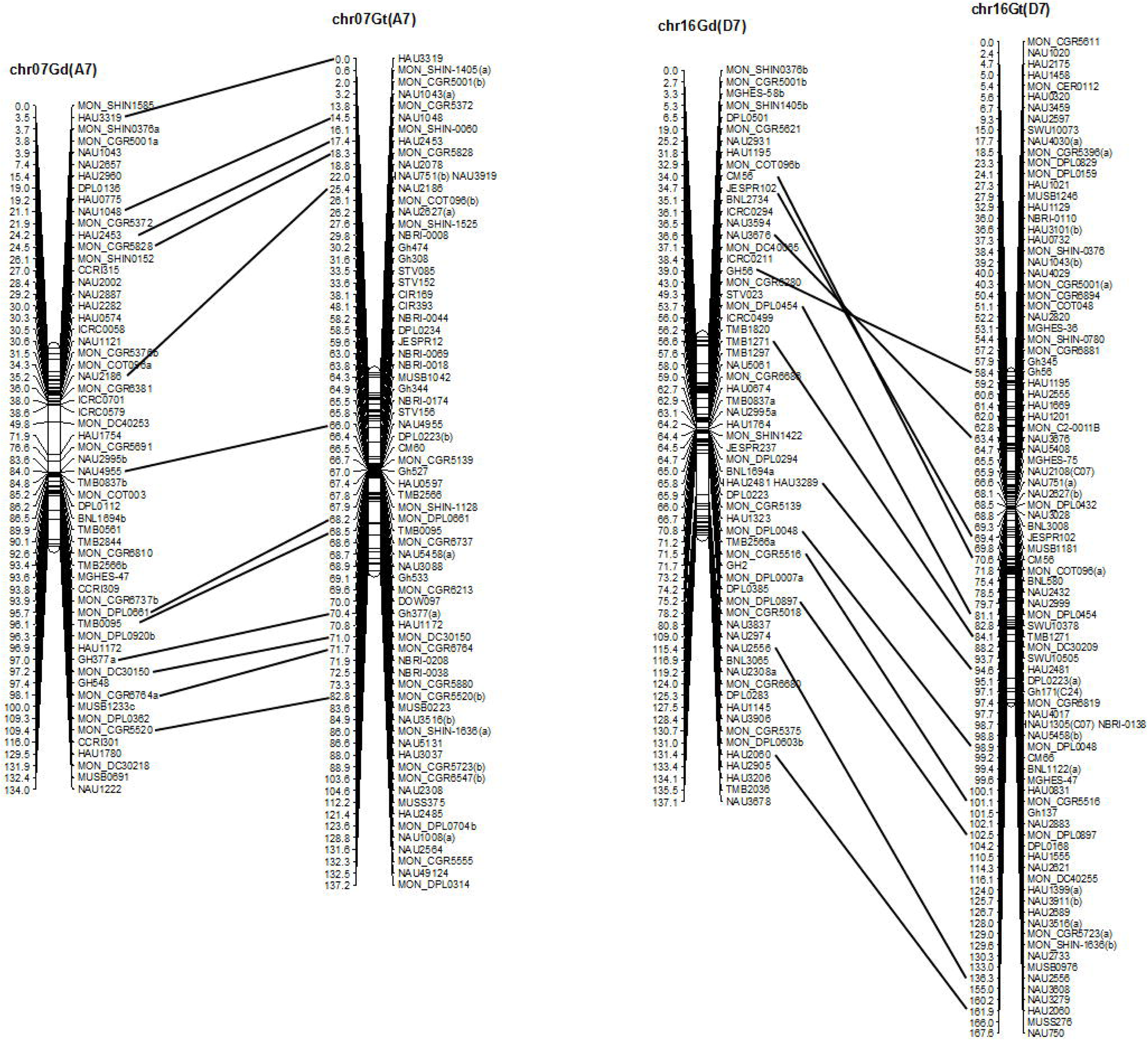

**Figure 8.**
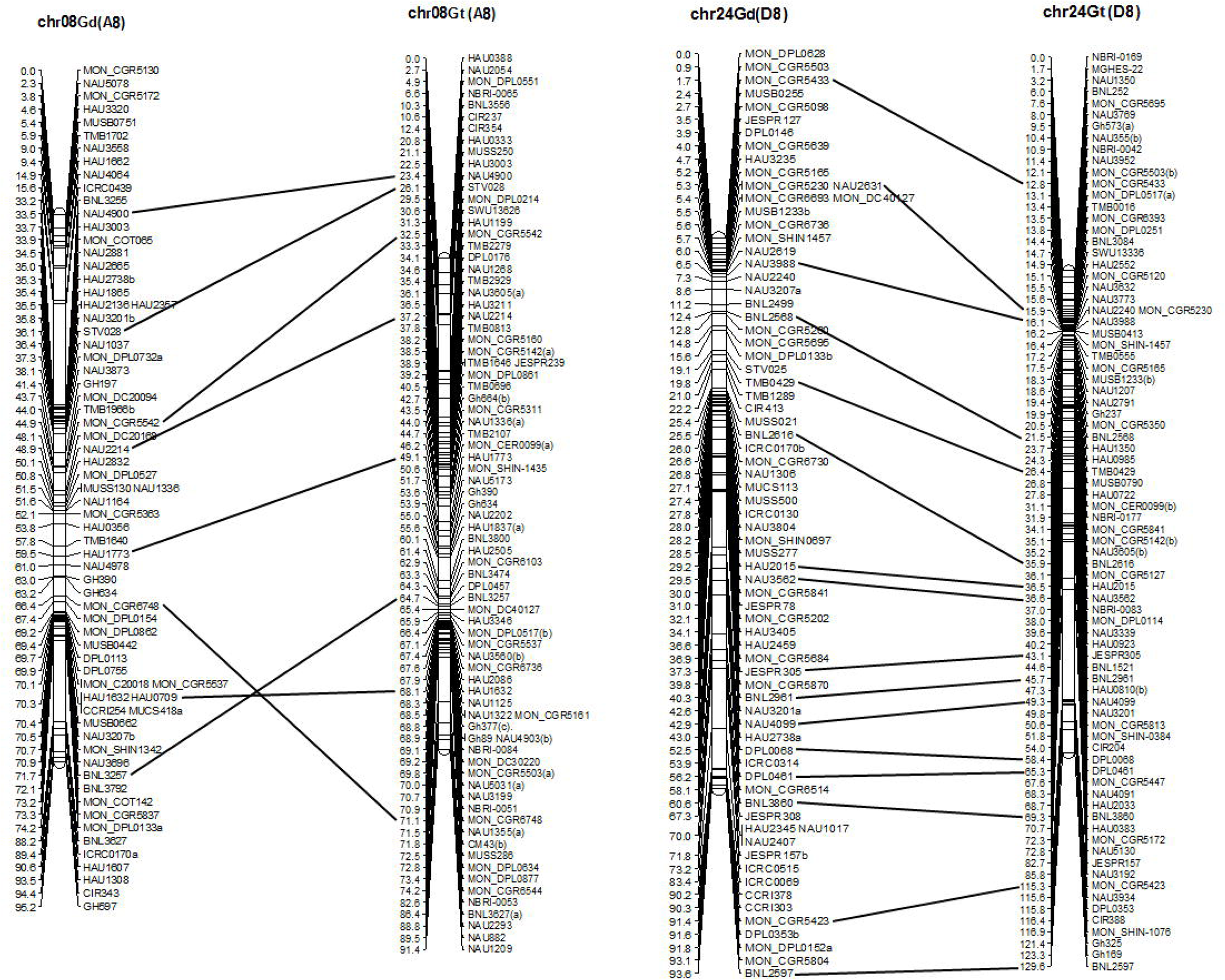

**Figure 9.**
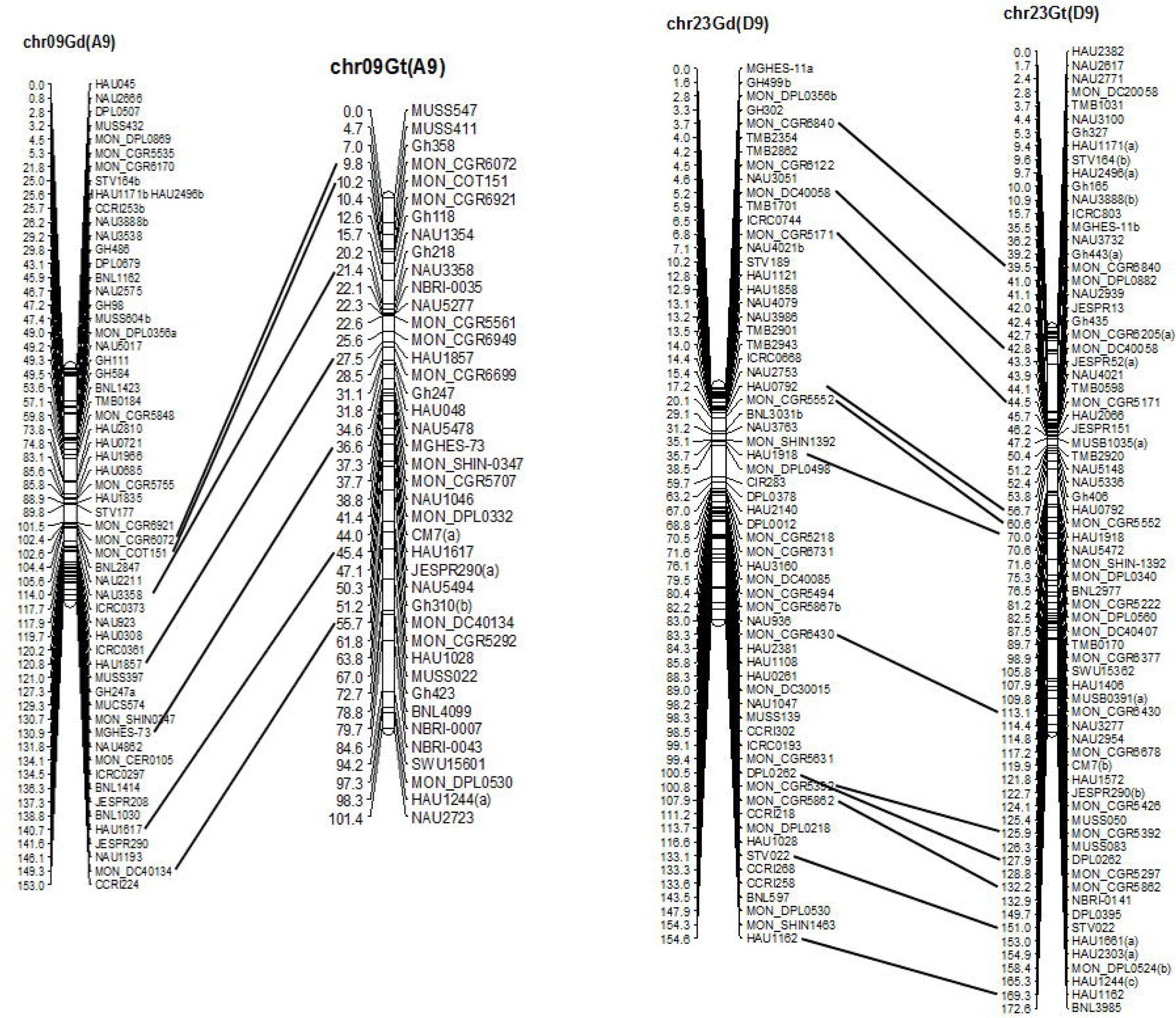

**Figure 10.**
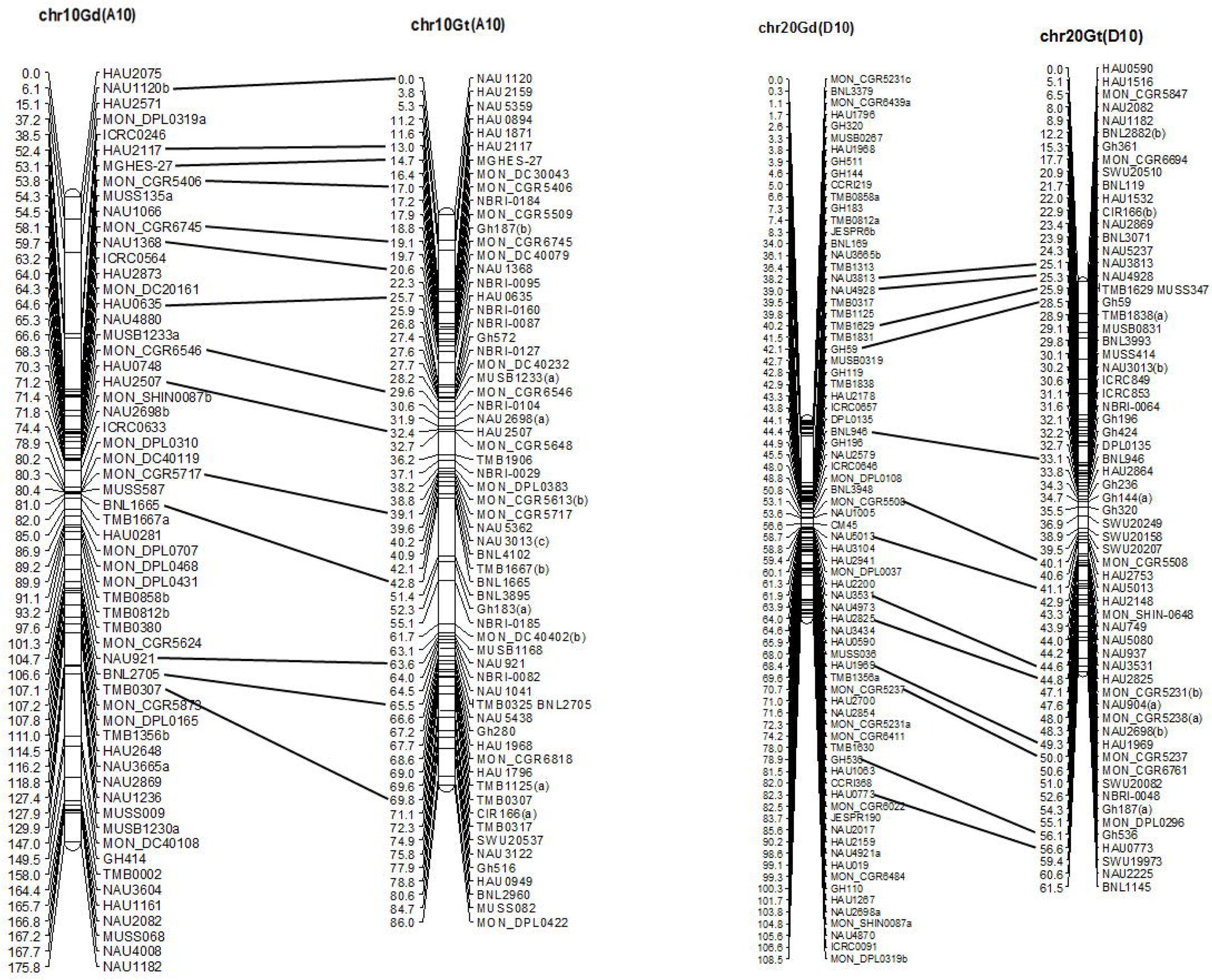

**Figure 11.**
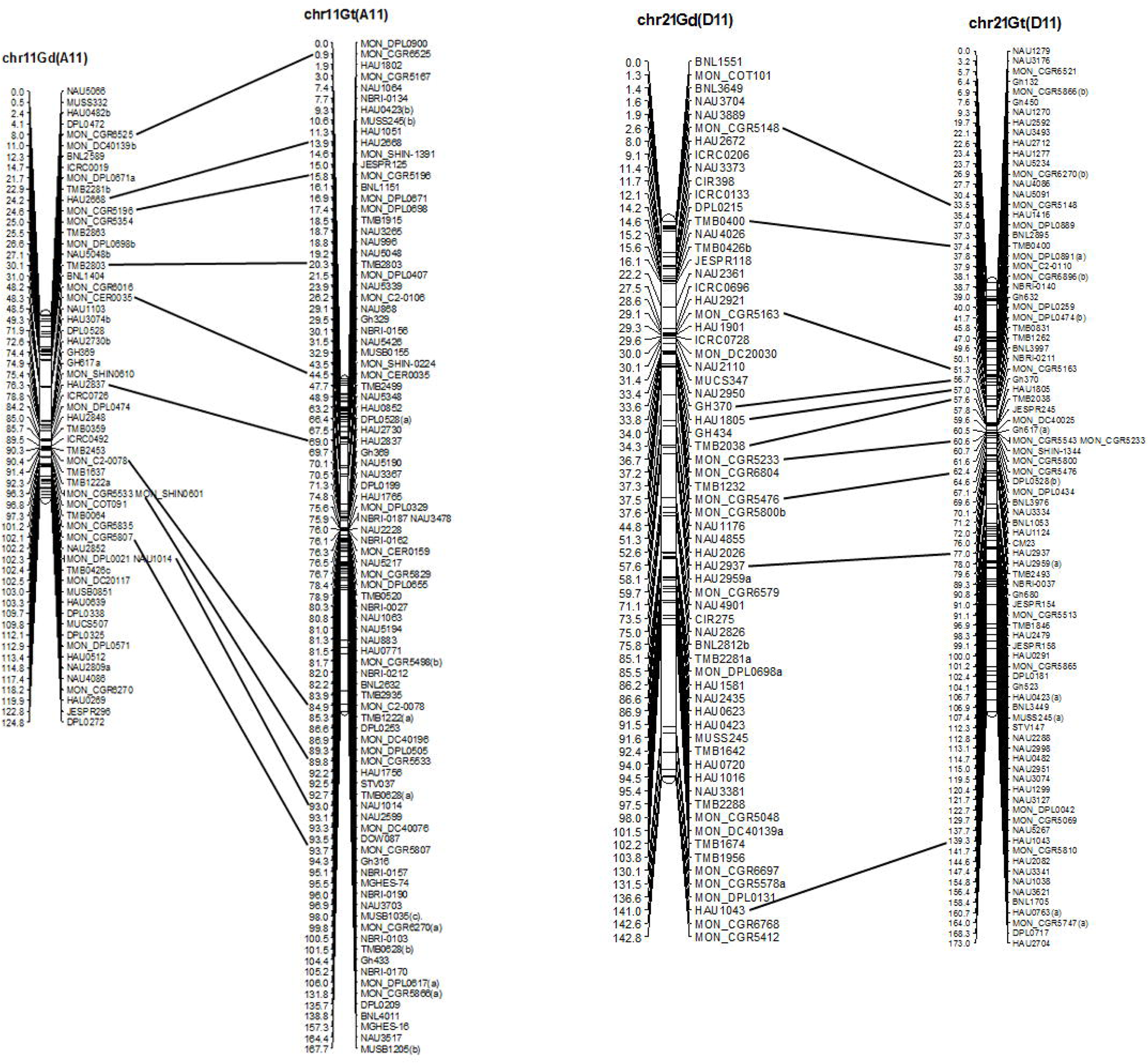

**Figure 12.**
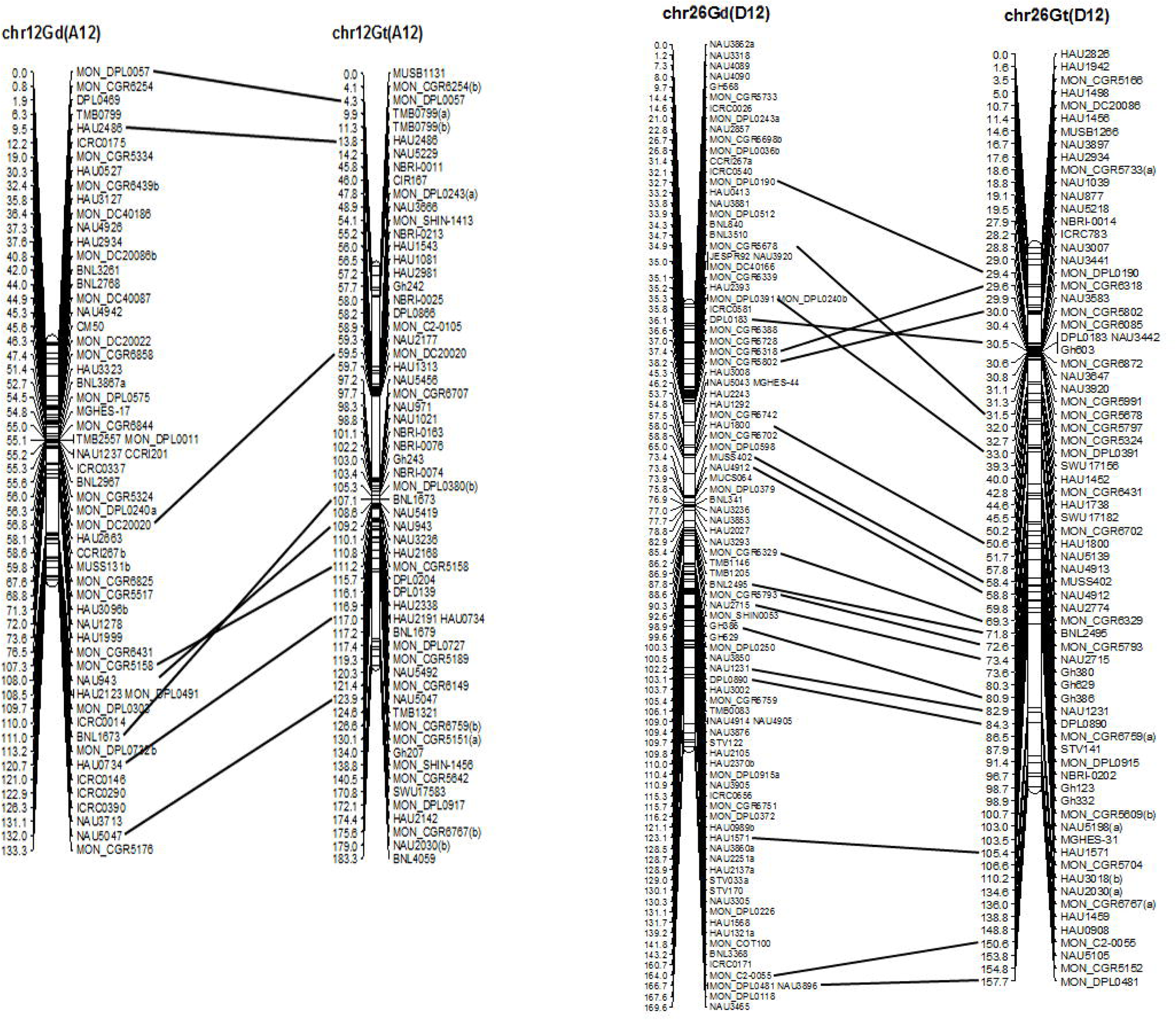

**Figure 13.**
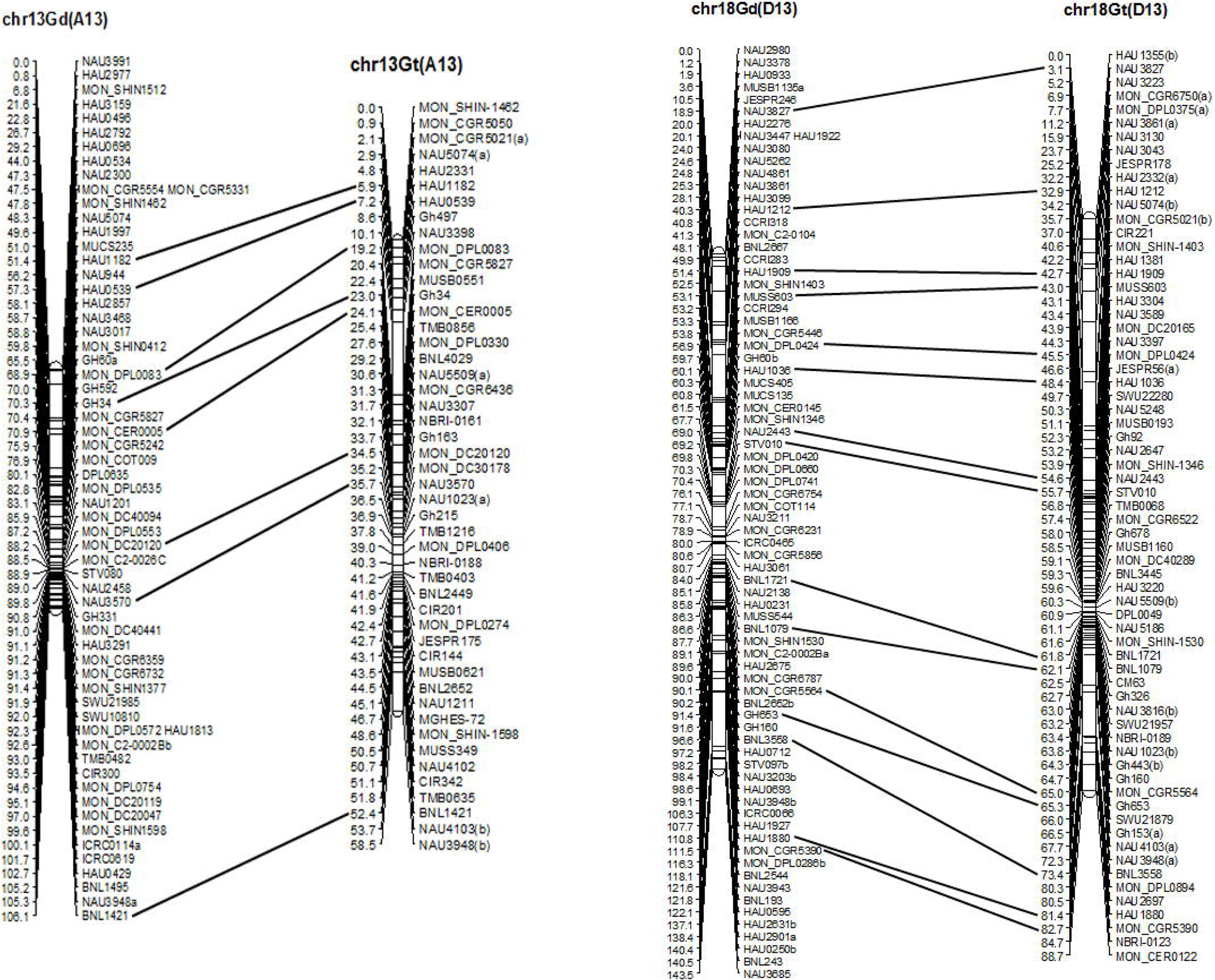

Seven inversions were found on senve different chromosomes between the genetic maps of *G. tomentosum* and *G. darwinii* respectively: on chr02 of MON_C0T043-HAU2643-HAU0248-HAU2923, on chr05 of MON-DPL0384-TMB1586-BNL2448, on chr08 of CGR6748-HAU0709-BNL3257, on chr12 of MON_CGR5158-NAU943-BNL1673, on chr14 of NAU3691-MON_CGR5818-TMB1513, on chr16 of CM56-JESPR102-NAU3676-GH56 and on chr25 of MON_CGR5665-NAU2388-BNL3103. Three translocations were found on 3 different chromosomes between the genetic maps of *G. tomentosum* and *G. darwinii* respectively: on chr05 of NAU2001-NAU1200-NAU3569 fragment parallel translocation with MUSB0592-NAU1127 fragment, on chr26 of MON-CGR5678-MON-DPL0391 fragment parallel translocation with MON-CGR6318-MON-CGR5802 fragment and on chr14 TMB0803-NAU2987 fragment parallel translocation with MON_C2_0079-HAU0551 fragment.

Referring to the published *G*. *Raimondii* genome sequencing data (D_5_, Wang et al. 2012) and the latest *G. hirsutum* genome sequencing data (AD_1_, Zhang et al. 2015), we compared the inversions and translocations on the genetic linkage map with the physical position and arrangement direction on the corresponding genome chromosome. The result showed that, among the inversions, the fragment orientation of *G. darwinii* on chr05 is consistent with the physical direction of *G. Raimondii* (D_5_) and *G. hirsutum* (AD_1_) genome sequence. The fragments orientation of *G. tomentosum* on chr08, chr12 and chr25 are consistent with the physical directions of *G. Raimondii* (D_5_). The fragments orientation of *G. tomentosum* on chr08, chr12 and chr16 are consistent with the physical directions of *G. hirsutum* (AD_1_). Among the translocation, just the fragment on chr26 corresponding physical position was found on *G. Raimondii* (D_5_), with the same direction as *G. tomentosum*.

## Discussion

### Reciprocal translocations and Inversion

In the study of *Gossypium* genetic linkage map construction, Researchers have repeatedly found in the reciprocal translocation between homologous chromosomes (Rong et al. 2004; He et al. 2007; Wang et al. 2006; John et al. 2012). Very few study focus on different cotton linear relationship between homologous chromosomes. In this study, use the same variety of *G. hirsutum,* we constructed allotetrapliod genetic linkage maps of *G. hirsutum* × *G. tomentosum* and *G. hirsutum* × *G. darwinii.* Through linear relationship analysis on homologous position, there are seven inversions (corresponding on chr02, chr05, chr08, chr12, chr14, chr16 and chr25) and three translocations (corresponding on chr05, chr14 and chr26) were found seperately on the two genetic linkage maps. Compare the inversions and translocations location with the physical position of *G. hirsutum,* we found the same inversion phenomena appeared on the corresponding position of chr02, chr05, chr12 and chr14 in *G. hirsutum,* the same with translocation on the corresponding position of chr05 in *G. hirsutum*. For the first time we discovered inversion on the homologous chromosome fragments of chr02 and chr14, and for the first time we found out the translocations on chr05 and chr14. In this study we also found inversion and translocation appeared at the same fragment, which indicate it is genetic active region. Further research need to confirm inversion occur before or after translocation.

### Genetic mapping coupled with physical alignment of genomic

A lot of research has been done on the origin and evolution of allotetraploid cotton by previous researchers. Fryxell (1992) and Wendel (2009) support the idea that *G. tomentosum* and *G. hirsutum* has a closer relationship, however *G. darwinii* and *G. barbadense* has a closer relationship. Rarely research has done to the linear relationship of chromosome genetic linkage between *G. tomentosum* and *G. darwinii.* With the cotton genome sequencing successively completed, relationship between genetic linkage map and physical map study became feasible and convenient. Using F_2_ population derived from interspecific crosses, we constructed two linkage maps of allotetraploid cotton *G. hirsutum* × *G. tomentosum* and *G. hirsutum* × *G. darwinii* separately. Referring to the latest published *G. hirsutum* genome sequencing data (Zhang et al. 2015), linear relationship of the two wild allotetraploid cotton genetic linkage maps was compared based on genome-wide SSR markers. The results show that the genetic linkage maps of *G. hirsutum* × *G. tomentosum* and *G. hirsutum* × *G. darwinii* had good linear relationship. Meanwhile, part of the non linear relationship appeared on the individual chromosomes between the two genetic linkage maps, including 7 inversion fragments and 3 translocation fragments. Compared with the sequence data of *G. hirsutum* (Zhang et al. 2015), we noticed among the 7 inversion fragments, just one markers orientation (chr05) which comes from *G. darwinii* has the same direction of physical position arrangement of *G. hirsutum,* while four markers orientation (chr08, chr12, chr16 and chr25) which comes from *G. tomentosum* has the same direction of physical position arrangement of *G. hirsutum or G. raimondii.* Among the 3 translocation fragments, one marker orientation (chr26) which comes from *G. tomentosum* has the same direction of physical position arrangement of *G. raimondii.* The result indicate compare *G. darwinii, G. tomentosum* has closer genetic relationship to *G. hirsutum.*

## Acknowledgements

This work was co-supported by the National Science and Technology Support Plan of China (2013AA102601) and National Key Technology Support Program of China (2013BAD01B03).

## Competing interests

The authors declare that they have no competing interests.

## Authors’ contributions

Kunbo Wang, Zhengsheng Zhang and Fang Liu participated in the experiments design. Fang Liu carried out most of the experiments and all computational analyses. Zhong-Li Zhou, Chun-Ying Wang and Yu-Hong Wang participated in part of groups construction, Xiao-Yan Cai, Xing-Xing Wang participated in part of mapping experiments. Fang Liu drafted the manuscript. Baohong Zhang, Kunbo Wang and Zhengsheng Zhang revised the manuscript. All authors read and approved the final manuscript.

---

Figure 1 to 13: genetic map and Collinearity comparing of *G. Tomentosum* and *G. Darwinii.*

Gd: *Gossypium Darwinii*

GT: *Gossypium Tomentosum*

**Table 1.** Location comparing of genetic linkage fraction and genome physical position

| Linkage group | Markers | Location on *G. darwinii*(genetic position, cM) | Location on *G. tomentosum*(genetic position) | Corresponding location on *G. raimondii* (physical position, Mb) | Corresponding location on *G. hirsutum* (physical position, Mb) | Events |
| --- | --- | --- | --- | --- | --- | --- |
| chr02 | MON_COT043-HAU2643-HAU0248-HAU2923 | 77.7-78.8-78.9-79.4 | 92.5-91.3-89.7-87.8 |  |  | inversion |
| chr05 | MON_DPL0384-TMB1586-BNL2448 | 144.1-146.1-147.8 | 86.3-84-82.6 | 21.4-21.6 | 37.2-58.2 | inversion |
| chr08 | MON_CGR6748-HAU1632-BNL3257 | 66.4-70.3-71.7 | 71.1-68.1-64.7 | 6.5-31.3-46.1 | 85.6-58.9 | inversion |
| chr12 | MON_CGR5158-NAU943-BNL1673 | 107.3-108-111 | 111.2-109.2-107.1 | 44.2-42.6 | 93.5-41.7 | inversion |
| chr16 | CM56-JESPR102-NAU3676-GH56 | 34-34.7-36.6-39 | 70.6-69.4-63.4-58.4 |  | 47.1-39.5 | inversion |
| chr25 | MON_CGR5665-NAU2388-BNL3103 | 18.6-20.9-23.3 | 29.8-29.4-28.6 | 42.9-40.6 |  | inversion |
| chr05 | NAU2001-NAU1200-NAU3569-MUSB0592-NAU1127 | 195.1-199.5-202.9-205.7-207 | 132.7-133.4-133.7-129-132.5 |  |  | translocation |
| chr26 | MON_CGR5678-MON_DPL0391-MON_CGR6318-MON_CGR5802 | 34.9-35.3-37.4-38.2 | 31.5-33-29.6-30 | 17.1-31.1-10.2-10.8 |  | translocation |
| chr14 | TMB0803-NAU2987-MON_C2_0079-HAU0551-NAU3691-MON_CGR5818-TMB1513 | 51.9-52.1-53.9-54.8-55-55.3-56.6 | 68-69.5-65.9-66.2-64.5-62.3-60.4 |  |  | inversion & translocation |

